# Automated and quantitative characterization of multi-scale benthic habitat and associated biological communities of an unknown southeast Pacific seamount

**DOI:** 10.64898/2026.03.11.710978

**Authors:** Yakufu Niyazi, Denise J. B. Swanborn, Jan M. Tapia-Guerra, Javier Sellanes, Erin E. Easton, Germán Zapata-Hernández, Heather A. Stewart, Alan J. Jamieson

## Abstract

Seamounts are prominent deep-ocean features that strongly influence geological processes, ocean circulation, and benthic biodiversity. Despite their importance, most seamounts remain unmapped and poorly characterized, particularly in the southeast Pacific Ocean, a region recognized for high marine endemism and ecological isolation. In this study, we present a quantitative habitat characterization of a previously undocumented seamount, informally named Solito Seamount, located between the Nazca–Desventuradas Marine Park and the Juan Fernández Archipelago. High-resolution multibeam bathymetry and backscatter intensity data were integrated with *in situ* observations from two remotely operated vehicle (ROV) dives (SO643 and SO645) to investigate how geomorphology and substrate distribution influence benthic community patterns. An automated and hierarchical quantitative mapping framework incorporating objective terrain analysis and multivariate statistical techniques, including principal component analysis and clustering, was applied to delineate five distinct megahabitat types: flat, basal slope, valley, ridge slope, and ridge crest. ROV video transects traversing these megahabitats revealed five associated substrate type forming macrohabitats: bedrock, bedrock with sediment veneer, sediment-rock transition, sediment, and coral rubble. Outputs were used to investigate how environmental heterogeneity structures megafaunal assemblages of Solito Seamount. Multivariate analysis revealed a combined effect of megahabitat type and substrate type on benthic megafaunal assemblages across the depth gradient. These compositional dissimilarities were primarily driven by habitat-forming taxa. In the deeper dive (SO643), a broad suite of taxa contributed to dissimilarities, and assemblages were primarily organised by megahabitat. The ridge crest hosted a distinct reef-building scleractinian community, whereas the ridge slope hosted mixed antipatharian, gorgonian and actiniarian assemblages. In contrast, the shallower dive exhibited simpler patterns with few taxa driving dissimilarities. Substrate effects were most pronounced with coral rubble forming a distinct habitat characterised by sponges (*Stelletta* sp.). Pronounced biological differences between dives may also represent depth-dependent structuring resulting from differences in oxygen regimes associated with water masses, underscoring the role of oceanographic forcing. This study provides the first quantitative habitat map of this previously undocumented seamount, delivering essential baseline information for this largely unexplored region of the southeast Pacific. The integrated multi-scale geophysical and biological approach presented here offers a robust framework for advancing seamount ecosystem understanding and supporting future biodiversity assessments and conservation planning.

## 1. Introduction

Seamounts are prominent bathymetric features of the deep ocean, rising more than 1000 m above the surrounding seafloor, exhibiting substantial variation in depth, morphological characteristics, and environmental setting [1–4]. These undersea mountains, primarily formed by volcanic activity, influence oceanographic processes that shape benthic and pelagic biological communities [5–7]. Formation and subsequent evolution of seamounts establish new source-to-sink systems that modify local hydrodynamics and sediment-transport pathways [8–12]. Volcanic and hydrothermal activity associated with seamounts plays a key role in generating metal-rich deposits, including polymetallic nodules, hydrothermal sulphides, sulphates, elemental sulphur, and phosphate accumulations [13–15]. The rugged topography of seamounts disrupts prevailing currents that can lead to localized turbulence and upwelling, altering oceanographic processes by redistributing particles and modifying thermal gradients [16–18].

Beyond their geologic and oceanographic significance, seamounts are widely recognized as ecological hotspots in the deep-sea, supporting high biological productivity, a diverse range of communities and endemism [19–21]. The geomorphology of seamounts, including variations in slope, relief, and substrate type, and the overlying hydrodynamic regime creates localized environmental conditions that play fundamental roles in structuring seafloor habitats and biological communities [22–24]. Flat or gently sloping summit plateaus, such as those on the Great Meteor Seamount in the North Atlantic, provide stable substrates that support mass accumulation of free-living scleractinians (stony corals) on soft sediment habitats [25]. In contrast, steep volcanic flanks enhance bottom currents, restrict sediment accumulation, and promote dense assemblages of suspension feeders as documented at the New England Seamount Chain, where slope, sediment types, and dissolved oxygen are the main factors explaining the distribution of diverse sponge taxa [26]. Observations along the Oahu Island of the Hawaiian Ridge suggested that the depth gradient is critical in structuring benthic communities, with small-scale physiographic variations such as the rugosity and slope playing a minor role [27]. The global distribution of seamounts and their three-dimensional complexity result in hotspots of biodiversity that further provide unreplaceable ecological connectivity [21,28]. Habitat heterogeneity, in turn, drives species distribution, community composition, and ecological interactions, contributing to the high benthic and pelagic diversity observed on seamounts [20,29–32].

Despite their important roles in facilitating enriched ocean biodiversity, only around 4% of global seamounts have been sampled for scientific purposes, with much of our understanding derived from a relatively small number of well-studied sites [33]. Therefore, many seamounts remain unmapped, and quantitative habitat characterization is particularly limited in the southeast Pacific Ocean, where only about one-fifth of the abyssal seafloor has been mapped at high resolution [34]. This lack of systematic exploration poses major challenges for biodiversity assessments, ecosystem modelling, and conservation planning in this region, which hosts numerous seamount chains formed by the Nazca (NZ), Salas y Gómez (S&G), and the Juan Fernández (JF) ridges. This region hosts exceptionally distinctive biological communities, including some of the highest levels of marine endemism recorded globally, and the deepest known light-dependent coral reefs, reflecting a combination of long-term ecological isolation and unique environmental conditions [35–38]. Most seamounts in this region occur along the NZ and S&G ridges, yet only around 3% have been scientifically studied since the 1970s, when expeditions by the former USSR reported high levels of endemism for benthic communities [39–41]. Several scientific campaigns have been conducted in the region, but they have focused almost exclusively on a limited number of ridge-associated features, leaving the vast majority of seamounts poorly characterized or completely unexplored [35,42–44]. To better understand the biodiversity, geological structure, and habitat complexity of these seamounts, several expeditions have been undertaken since 2024. One of which is the Schmidt Ocean Institute’s 2024 expedition FKt240108 (Seamounts of the Southeast Pacific, https://schmidtocean.org/cruise/seamounts-of-the-southeast-pacific/) on board the R/V *Falkor (too)*. During this campaign, an undocumented and formally unnamed seamount (Solito, Fig. 1) located between the Nazca-Desventuradas Marine Park and the Juan Fernández Archipelago was investigated. In this study, we use an automated and quantitative mapping approach to describe the habitat characteristics of this newly mapped seamount and to investigate how different megahabitats and substrate types drive the spatial patterns of benthic communities. Quantitative mapping of seafloor habitats combines geophysical and biological techniques to systematically describe the seafloor and their associated communities [45–50].

**Figure 1.**
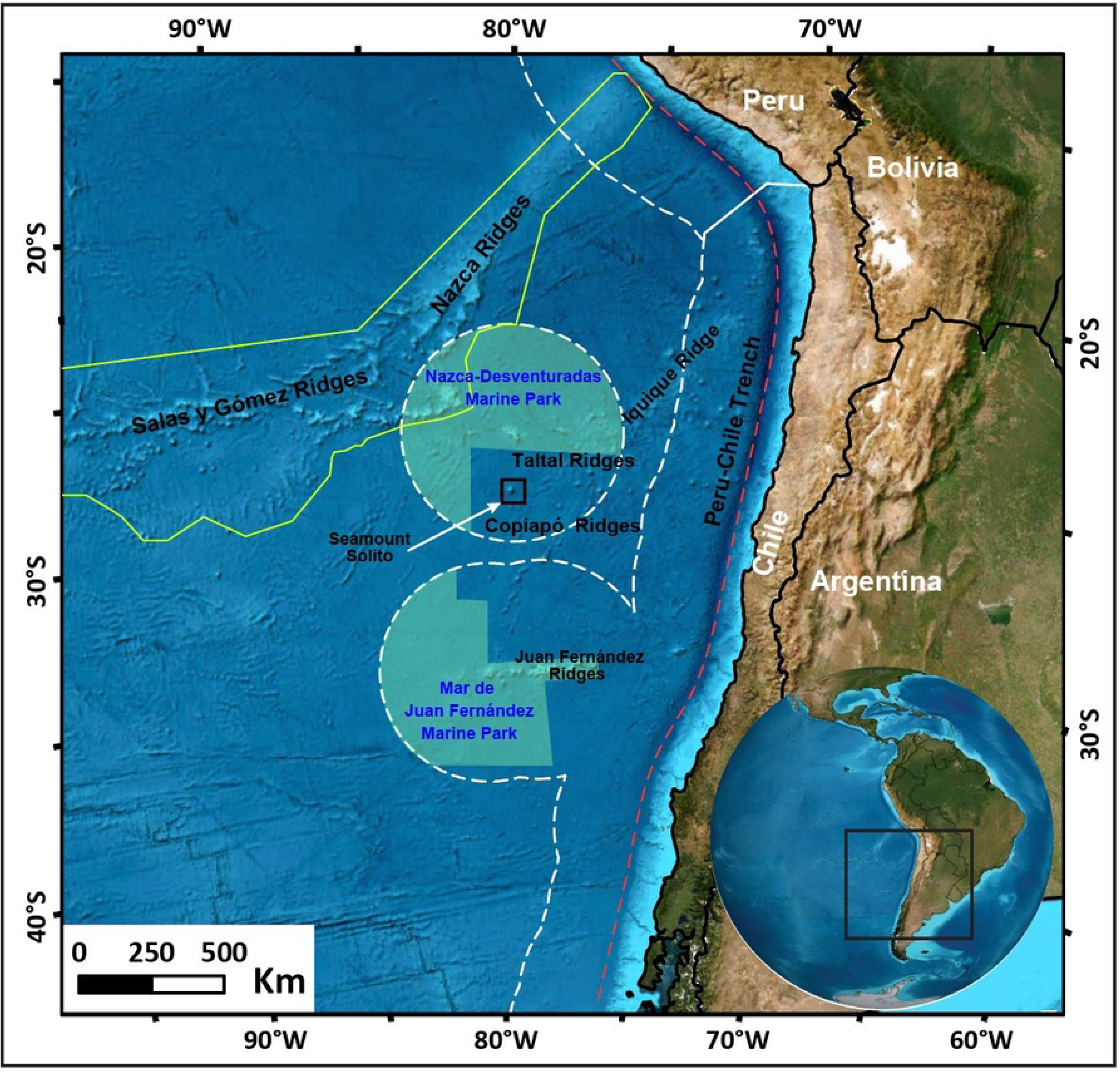
Overview map of the study area located in the southeast Pacific Ocean. The informally named Solito Seamount, the focus for expedition FKt240108, is indicated by the black box. The white dashed line represents the Exclusive Economic Zone. Green polygonal areas correspond to marine parks, and the green line outlines the Nazca and Salas & Gómez Ecologically or Biologically Significant Marine Area (EBSA) designated by the Convention on Biological Diversity. Red dashed line indicates the subduction zone between the Pacific and South American tectonic plates. Bathymetry is from the General Bathymetric Chart of the Oceans (GEBCO) (GEBCO Compilation Group, 2025).

A critical component of seafloor habitat characterization is the application of hierarchical analytical frameworks capable of capturing the multiple spatial scales that influence biodiversity patterns [50–52]. Seamounts exhibit strong geomorphological heterogeneity, with geological and oceanographic processes operating across multiple spatial scales, from mega-scale physical seafloor habitats to macro-scale substrate types and benthic community patterns, offering insights into the processes shaping deep-sea ecosystems [6,53–55]. Yet, automated and quantitative multiscale habitat assessments remain relatively rare in deep-sea environments, largely due to limitations in data resolution, coverage, and standardization. Several approaches exist for habitat classification, ranging from semi-automated to fully automated methods. Habitat classification using the Benthic Terrain Modeler (BTM) typically relies on user-defined thresholds that can reduce consistency and reproducibility across studies [56–59]. Similarly, rule-based classifications of bathymetry and backscatter rely heavily on expert judgement, requiring iterative testing and introducing subjectivity into habitat assignment [60–62]. While these approaches facilitate terrain analysis and seabed segmentation, they are inherently constrained by expert bias and limited reproducibility.

Automated and quantitative characterization of multi-scale benthic habitat, by contrast, offers an objective means of integrating multiple terrain and acoustic predictors to delineate ecologically meaningful megahabitats [63–66]. In this study, the combined use of terrain derivatives, principal component analysis, and unsupervised clustering enables dimensionality reduction and the objective identification of megahabitat classes across the seamount. *In situ* observations from remotely operated vehicles (ROVs) allow identification of substrate types and direct assessment of benthic assemblages, enabling explicit links between megahabitat, substrate characteristics, and species composition and distributions. This hierarchical approach captures the multi-scale nature of seamount ecosystems, from mega-scale geomorphology to macro-scale substrate mosaics and biological communities, providing mechanistic insight into the processes shaping benthic biodiversity. As such, the automated and quantitative framework applied here offers a reproducible, scalable, and ecologically interpretable approach for seamount habitat characterization, supporting robust ecosystem assessments and informing conservation planning in data-poor regions of the deep-sea.

## 2. Materials and methods

### 2.1 Study area

This study focuses on a remote submarine feature, unofficially named Solito Seamount, located in the southeastern Pacific Ocean within Chile’s Exclusive Economic Zone. The seamount lies outside of the Nazca-Desventuradas Marine Park, roughly 200 km south of the Desventuradas Islands and roughly 620 km north of the Juan Fernández Archipelago (Fig. 1). Despite its prominence, the tectonic and volcanic origin of Solito remains unresolved. One hypothesis links it to the Iquique Ridge that is currently being subducted beneath the South American Plate [67]. Alternatively, Solito Seamount may represent the westernmost extension of the east–west trending Copiapó Ridge that runs subparallel to the Taltal and JF ridges and is also thought to be subducting beneath the adjacent continental margin [68]. However, the limited availability of geochronological and geochemical data for these ridges and Solito constrains efforts to reconstruct their geological evolution.

Oceanographically, the southeastern Pacific is structured by vertically stratified water masses that regulate temperature, oxygenation, and near-bottom circulation [69,70]. Surface circulation includes the Humboldt Current and associated coastal branches, supporting high primary productivity along the Chilean coast and continental margin [71,72]. Below surface waters, a persistent oxygen-depleted and nutrient-enriched Equatorial subsurface water (ESSW, ∼50-600 m) is associated with the regional Oxygen Minimum Zone (OMZ). At intermediate depths (600–1,300 m), well-oxygenated Antarctic Intermediate Water (AAIW) flows northward, whereas nutrient-rich and well-oxygenated Pacific Deep Water (PDW; ∼1,300–3,000 m) occupies deeper levels. Below the PDW, Circumpolar Deep Water (CDW), together with contributions from Antarctic Bottom Water (AABW) and North Atlantic Deep Water (NADW), dominates abyssal circulation [72]. From a biogeographic perspective, Solito occupies a strategic position between the NZ, S&G, and JF ridges, that may form part of a proposed connectivity corridor that may enable faunal exchange between isolated oceanic islands and continental deep-sea ecosystems [35,37,43,44].

### 2.2 Data collection

#### 2.2.1 Multibeam echosounder data

The expedition aimed to produce detailed seafloor maps alongside direct geological and biological observations. High-resolution bathymetry and acoustic backscatter intensity measurements were collected using a hull-mounted Kongsberg EM 124 multibeam echosounder (MBES). Operating at a nominal frequency of 12 kHz within a 10.5–13.5 kHz range, the system has a beam width of 0.5° × 1° and is optimized for depths between 1,000 and over 8,000 m. Data acquisition was conducted via Kongsberg’s Seafloor Information System (SIS, version 5) with vessel positioning supplied by a Seapath 380+ navigation system. Sound velocity profiles were recorded using a Reson SVP70 sensor attached to the transducer head, complemented, when necessary, by expendable bathythermograph (XBT) drops. All raw data were processed in QPS Qimera software suite (version 2.5) with final bathymetric digital elevation models (DEMs) gridded and exported at a resolution of 100 m pixel size (Fig. 2). The FMGeocoder Toolbox (version 7.10) for Fledermaus was used to produce backscatter intensity mosaics at a resolution of 50 m pixel size, depicting acoustic reflectivity patterns.

**Figure 2.**
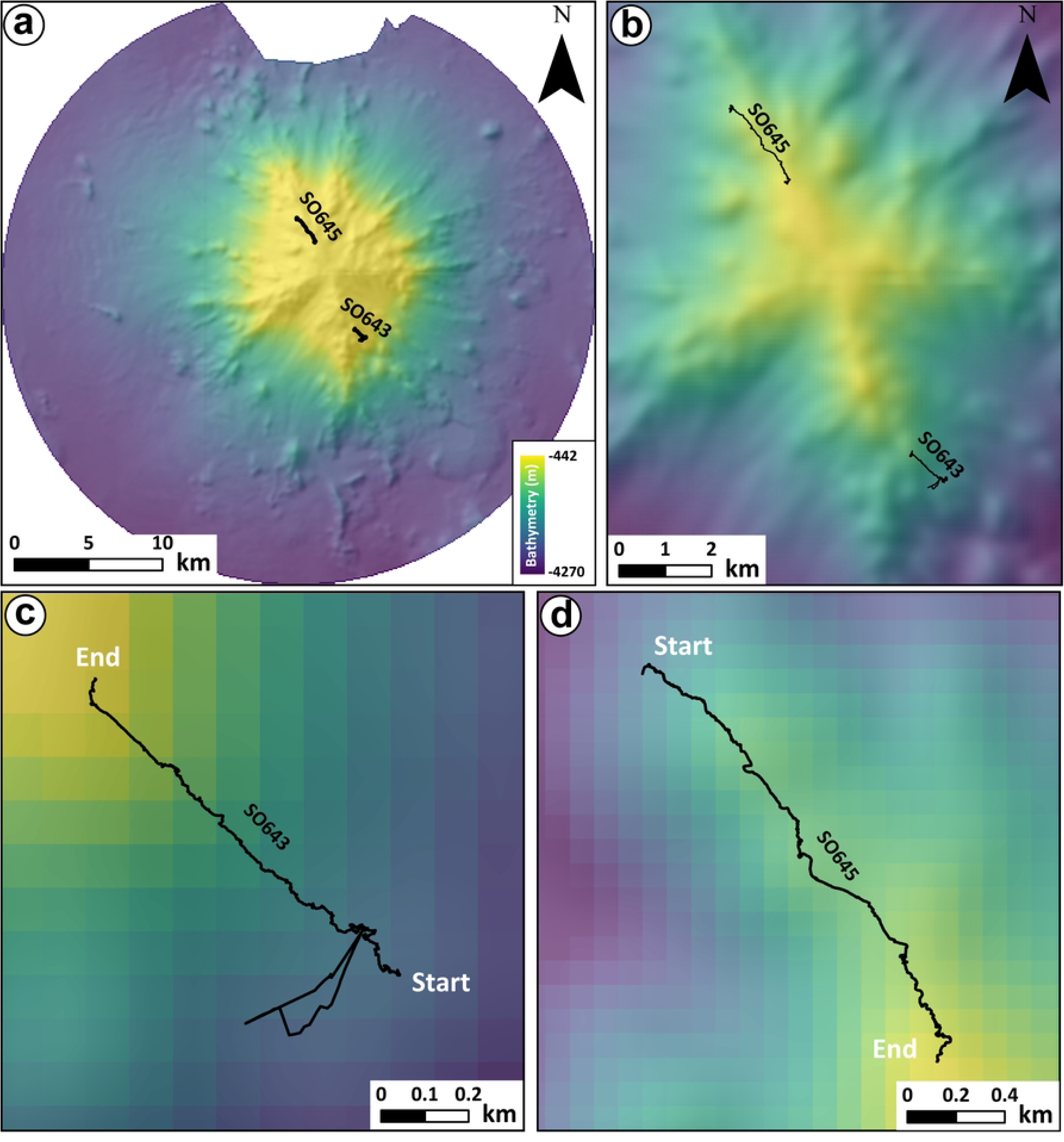
(a) High-resolution multibeam bathymetry of the Solito Seamount and remotely operated vehicle (ROV) dive tracks (black lines) acquired during expedition FKt240108. (b) Zoom in of the seamount summit and ROV dive locations. Close up ROV dive tracks for (c) SO643 and (d) SO645.

#### 2.2.2 ROV data

The 4,500-m–rated ROV *SuBastian* conducted benthic video surveys using high-definition (HD) and 4K ultra-high-definition (UHD) cameras. The ROV is equipped with two 4K Sulis Subsea Z70 (resolution 840 × 2160 pixels) deep-sea video cameras and four DSPL FlexLink HD Multi SeaCams. Two ROV dives were undertaken on Solito at sites selected based on the location of topographic ridges and hardgrounds, interpretated from MBES data (Fig. 2). These dives were conducted on the southern flank at ∼1,054–1,497 m (lower to mid slope dive, SO643) and on the northern flank at ∼456–860 m (upper slope to summit dive, SO645) (Table 1, Fig. 2). The ROV transects primarily advanced upslope, surveying benthic invertebrates, benthopelagic organisms, and seafloor geology. The surveys yielded approximately 16 h of HD video footage for benthic surveys and specimen collections. Logging of observations and sampling events were made using the Sealog annotation system [73,74], which allowed systematic logging of frame grabs together with environmental and positional metadata. For benthic fauna assessments, frames were captured every 10 s from the videos using the ASNAP (auto snapshot) function in the Sealog software to ensure consistent and reliable frame extraction, providing accurate and unbiased sampling for subsequent abundance analyses.

**Table 1.**
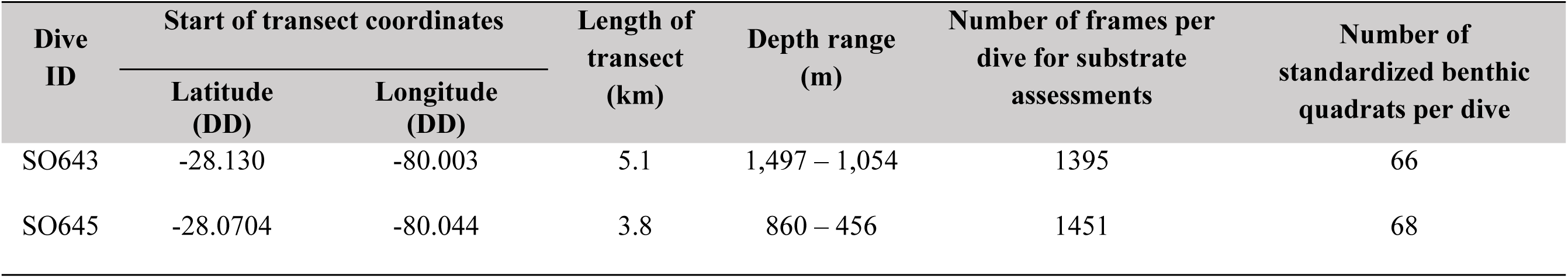
Summary of ROV dive operations conducted at the Solito Seamount during expedition FKt240108.

### 2.3 Data analysis

#### 2.3.1 Quantitative megahabitat mapping

To characterise mega-scale seafloor habitats (hereafter megahabitats) across Solito, we first applied quantitative terrain analysis using the multibeam bathymetry and backscatter intensity data gridded at 100 m and 50 m resolution respectively. A suite of seafloor terrain derivatives were calculated from the bathymetric grid to characterize geomorphological complexity. These derivatives include slope, bathymetric position index (BPI), profile curvature, planform curvature, terrain ruggedness index (TRI), and vector ruggedness measure (VRM) (Table S1). Slope and curvature metrics characterize local and along-slope seabed morphology, providing insights into the fine-scale structure of the seafloor. In contrast, BPI, TRI, and VRM quantify broader aspects of topographic complexity and surface heterogeneity. Collectively, these bathymetric derivatives have been extensively applied in both quantitative and qualitative habitat mapping and have demonstrated strong utility in describing the geomorphological characteristics of seamount ecosystems [48,49,56,66,75,76]. An internal radius of 100 m and an external radius of 300 m were used to calculate fine-scale BPI, while an internal radius of 200 m and an external radius of 500 m were adapted to calculate broad-scale BPI, thereby capturing variations across multiple scales (S1 Fig. 1). Backscatter intensity data were incorporated as a proxy for the distribution of soft and hard substrate on the seabed, providing information on sediment and hardground distribution [77–80].

Because the two ROV transects did not span the full depth range of Solito, they provided only limited representative training data for supervised classification. Therefore, a broad-scale habitat classification was performed using established unsupervised methods to identify natural groupings in the environmental variables [50,63,81]. Principal component analysis (PCA) was performed [82] to reduce data dimensionality and keep the most informative and uncorrelated components to capture the majority of variance in the dataset [83]. Variables were scaled and centred before conducting the PCA to ensure they had equal weight in the analysis. To classify seafloor habitats, k-means clustering was applied to the retained principal components (PCs) [84]. The optimum number of clusters was determined via an elbow plot. The characteristics of each cluster were evaluated using boxplots of the PCA scores [85]. The resulting clusters were mapped back onto the original bathymetric surface, producing a spatially explicit representation of benthic megahabitats as defined by broad-scale geomorphology [51,55,86–88]. All quantitative habitat mapping analyses were conducted using the open-source software R (version 4.4.2) [89] and the packages *raster*, *RStoolbox*, *terra*, *reshape2*, *MultiscaleDTM*, and *spatialEco* [90–92].

#### 2.3.2 Macro-scale substrate characterisation from ROV imagery

To provide macro-scale ground truthing, ROV video data were used to classify substrate types throughout the dives. Each ROV video transect was first reviewed in full to gain a general understanding of the substrate variability and morphological context. Then video frames were exported as still images at 30-second intervals for detailed annotation generating a total of 2,846 benthic still images for review. Images with low visibility (due to sediment or lighting) were excluded. Substrate types were classified according to the relative occurrence and predominance (>80%) of three main components: bedrock, sediment, and biogenic material. This classification was conducted through quantitative visual interpretation of seabed imagery, by comparing and adapting established substrate classification frameworks that are widely used in marine habitat studies [51,86,93–96]. Each image was assigned to one of the defined substrate classes with the predominant substrate composition, sediment cover, and structural complexity examined.

#### 2.3.3 Benthic fauna assessments

A subset of ROV ASNAP video frames were established as quadrats and were defined based on the following criteria: (i) a field of view between 1 and 5 m² to ensure accurate species identification and (ii) frames in which the ROV maintained a steady altitude between 1 and 5 m above the substrate, ensuring the visibility of the laser scalebar (two points positioned 10 cm apart) for precise measurement. Of these frames, a total of 134 frames, one per every ∼15 min of ROV dive time, were used for the standardized photographic quadrats (1–5 m²) [97 in review], comprising 66 from SO643 and 68 quadrats from SO645 (Table 1). Note that the dive transects were primarily focused on benthic (sessile and mobile) and benthopelagic organisms, thus fish were not considered in this research because no standardized method for fish abundance was used during data acquisition. Taxa were identified to the lowest possible taxonomic level based on regional taxonomic literature and previous faunal inventories from the Nazca–Desventuradas Marine Park and the Juan Fernández Archipelago [98–103], complemented by specimens collected during the present expedition. Specimens were collected using the ROV manipulator, preserved in 96% ethanol, and deposited in the Sala de Colecciones Biológicas of the Universidad Católica del Norte (SCBUCN) for detailed laboratory examination. Specimen collection and handling were conducted under permit R. EX. N° E-2023-612, issued by the Chilean National Fisheries and Aquaculture Service (SUBPESCA) to Universidad Católica del Norte, Chile. All identifications were subsequently cross validated using authoritative global databases, including the World Register of Marine Species (www.marinespecies.org) and the Ocean Biodiversity Information System (www.obis.org)

#### 2.3.4 Statistical analysis

A hierarchical framework was used to evaluate the structuring effect of multi-scale habitat characteristics on benthic biodiversity patterns at Solito. Differences in benthic assemblage structure between megahabitat classes and substrate types were first assessed using a Permutational Multivariate Analyses of Variance (PERMANOVA) in PRIMER version 7.0 on a Bray-Curtis dissimilarity matrix of square-root-transformed data and over 9,999 permutations. Dive was included as a random factor, megahabitat as a fixed factor nested in dive, and substrate as a fixed factor nested in megahabitat and dive. Follow-up pairwise tests were used to assess which megahabitat types and substrates significantly differed based on benthic assemblage structure.

To identify the taxa contributing most strongly to compositional differences among megahabitat and substrate types per dive, Similarity Percentage Analyses (SIMPER; [104]) were applied in R (version 4.4.2) [89] using package *vegan* [105] and *tidyverse* [106]. The dive community datasets were cleaned to exclude empty taxa (columns with zero counts), empty samples (rows with zero counts), and rare taxa (occurring in < 2 frames). SIMPER analyses were performed over 9,999 permutations on the cleaned community matrices using Bray-Curtis dissimilarity. SIMPER analyses were conducted separately for each dive to avoid confounding megahabitat-substrate contrasts with between-dive differences. Only megahabitat-substrate combinations represented by at least two replicate frames were retained for the analysis. SIMPER was performed for megahabitat (e.g. ridge crest and ridge slope), substrate (e.g. bedrock and sediment) as well as a combined megahabitat and substrate factor (e.g. ridge crest-bedrock and basal slope-sediment) per dive. SIMPER outputs were tidied and combined across dives by extracting the taxa contributing significantly (p <0.05) to group dissimilarities for each contrast. Taxa were assigned to these groups to identify taxon-habitat associations within each dive. Contributions were aggregated to create heatmaps of taxa most strongly associated with (1) each megahabitat grouping, (2) each substrate grouping, and (3) each combined megahabitat-substrate grouping.

## 3. Results

### 3.1 Seamount megahabitats

The PCA of the nine abiotic variables reveals clear groupings among geomorphological, terrain, and acoustic predictors (S1 Fig, S1 Table) with the five PCs explaining 93.9% of the total variance. The first two components explain a combined 67% of the total variance (38.4% and 28.6%, respectively), indicating they capture most of the essential structure in the data (S2 Fig). The inclusion of the third, fourth, and fifth components substantially increases the cumulative variance, indicating that these components represent additional but still meaningful gradients. The marked decline in explained variance beyond the fifth component suggests that higher-order components contribute little additional information and were therefore excluded from further analysis. The rotated component matrix shows the factor loads that explain the correlations between the rotated PCs and the original variables (S2 Table). Bathymetric position and curvature metrics (broad-BPI, fine-BPI, planform curvature, and profile curvature) load strongly on the first component, indicating a dominant geomorphological gradient separating crests, flanks, and depressions (Fig. 3, S2 Table). Slope and TRI are the primary contributors to the second component, reflecting gradients in steepness and relative topographic relief, whereas backscatter intensity overwhelmingly defines the third component, isolating a substrate reflectivity signal (S2 Table). VRM and curvature metrics contribute to higher-order components, capturing fine-scale surface complexity. Correlation patterns (Fig. 3) further indicate a strong negative relationship between depth and backscatter intensity, suggesting lower acoustic reflectivity in deeper areas, and a positive association between slope and TRI. Overall, the retained PCs summarise the major physical drivers of habitat heterogeneity on Solito Seamount and provide an ecologically interpretable, low-collinearity set of predictors for subsequent habitat mapping analyses.

**Figure 3.**
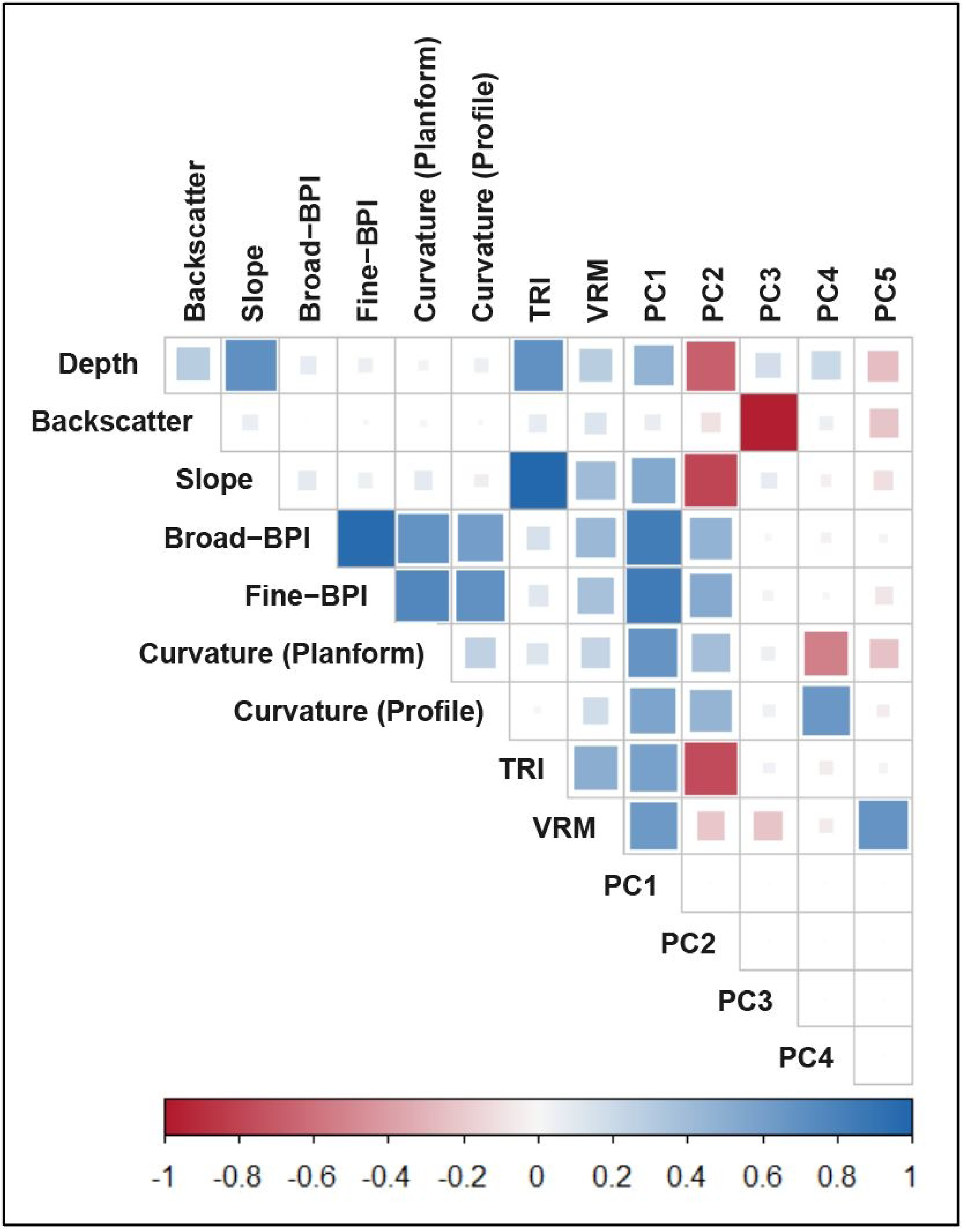
Correlation heat map of depth, backscatter intensity, slope, BPI, curvatures, TRI, VRM and main principal components. BBPI and FBPI: broad and fine-scale bathymetric position index; TRI: terrain ruggedness index, VRM: vector ruggedness measure.

The k-means clustering analysis, with five clusters selected using the elbow method, revealed distinct and spatially coherent megahabitat patterns across the seamount (Fig. 4). Violin–boxplots of standardized of geomorphometric and acoustic variables show clear differences among the five megahabitat classes, and allow these classes to be associated with five morphological structures of the seamount ([1] ridge crest, [2] ridge slope, [3] basal slope, [4] valley, and [5] flat seafloor) (Fig. 4, S3 Fig), indicating that the classification effectively captures the major variations in seafloor morphology and acoustic response. Megahabitat 1 is one of the most distinct megahabitats, interpreted as seamount ridge crests, characterized by relatively shallow depths, strongly positive BPIs, elevated TRI and VRM, and generally moderate to high slopes. These attributes are consistent with topographically elevated, structurally complex localized ridge crests (S3 Fig). Megahabitats 2 is also associated with elevated and structurally complex terrain but differs from ridge crest by occurring on steeper slopes with slightly greater depths and lower BPI values (S3 Fig). Megahabitat 3 (basal slope) occupies intermediate depths and is characterized by moderate slopes, slightly negative BPI values, and comparatively low ruggedness metrics, and is consistent with lower flank settings that form a geomorphic transition between steeper ridge slope and adjacent low-relief terrain (S3 Fig). Megahabitat 4 (valley) occurs at greater depths and is characterized by negative BPI values, gentle slopes, low backscatter intensity, and reduced terrain ruggedness, indicating depressed topographic settings that likely promote sediment accumulation and smoother substrate conditions (S3 Fig). Megahabitat 5 (flat seafloor) defines extensive low-relief areas surrounding the seamount. It is characterized by greater depths, minimal slopes, near-zero curvature, low TRI and VRM values, and generally low acoustic backscatter, consistent with smooth, laterally extensive abyssal to near-abyssal plain environments (S3 Fig).

**Figure 4.**
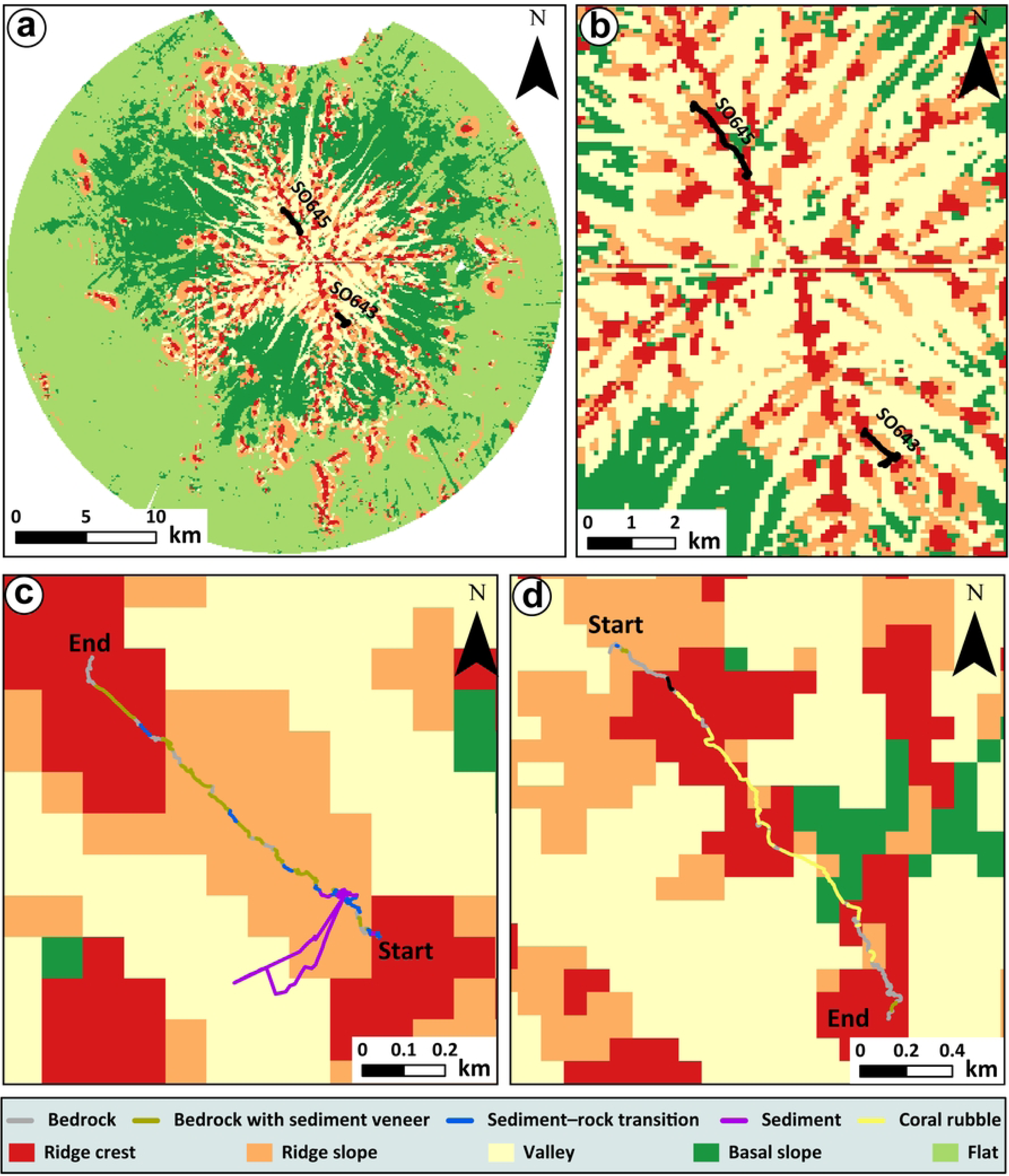
(a) Megahabitat classification for Solito Seamount and remotely operated vehicle (ROV) dive tracks (black lines). (b) Zoom in of the habitat classification for the seamount summit with ROV dive locations (black lines). Close up ROV dive tracks with observed substrate types for (c) dive SO643 and (d) dive SO645, overlain on habitat classification for the whole seamount.

### 3.2 Substrate types and distribution

Five substrate types were identified along the ROV transects: bedrock, bedrock with sediment veneer, sediment–rock transition, sediment, and coral rubble (Table 2; Figs. 4–6). Outcropping bedrock consists of exposed volcanic lava flows with rough surfaces, high structural complexity, and little to no (<5%) sediment cover. Bedrock with sediment veneer comprises volcanic bedrock partially draped by a thin (5–20%) layer of sediment (Table 2; Fig. 5). Sediment–rock transition substrates form heterogeneous mosaics of exposed bedrock, talus cobbles, and unconsolidated sediments, exhibiting moderate to high small-scale heterogeneity and variable sediment cover (10–50%). Sediment substrates are dominated by unconsolidated coarse-grained material forming smooth, low-relief seafloor with low structural complexity and high (>80%) sediment coverage. Coral rubble consists of accumulations of fragmented coral skeletons and skeletal debris mixed with sediment, displaying low to moderate structural complexity and variable (20–50%) sediment coverage. Both ROV dives crossed multiple seamount megahabitats and substrate types (Figs. 4 and 6). However, the overall substrate composition differed between the two transects. ROV dive SO643 showed relatively heterogeneous substrates (Figs. 4c and 6). Bedrock with sediment veneer accounts for roughly 60%, and exposed bedrock constituted approximately 20% of all frames observed in dive SO643 (Fig. 6c). While sediment–rock transition and sediment substrates each contributed about 10% of observations. In contrast, SO645 was strongly dominated by exposed bedrock, which comprised >80% of all frames (Fig. 6d). Coral rubble formed a minor component (∼12%), and bedrock with sediment veneer accounted for <10% of observations. Sediment and sediment–rock transition substrates were absent along this transect.

**Figure 5.**
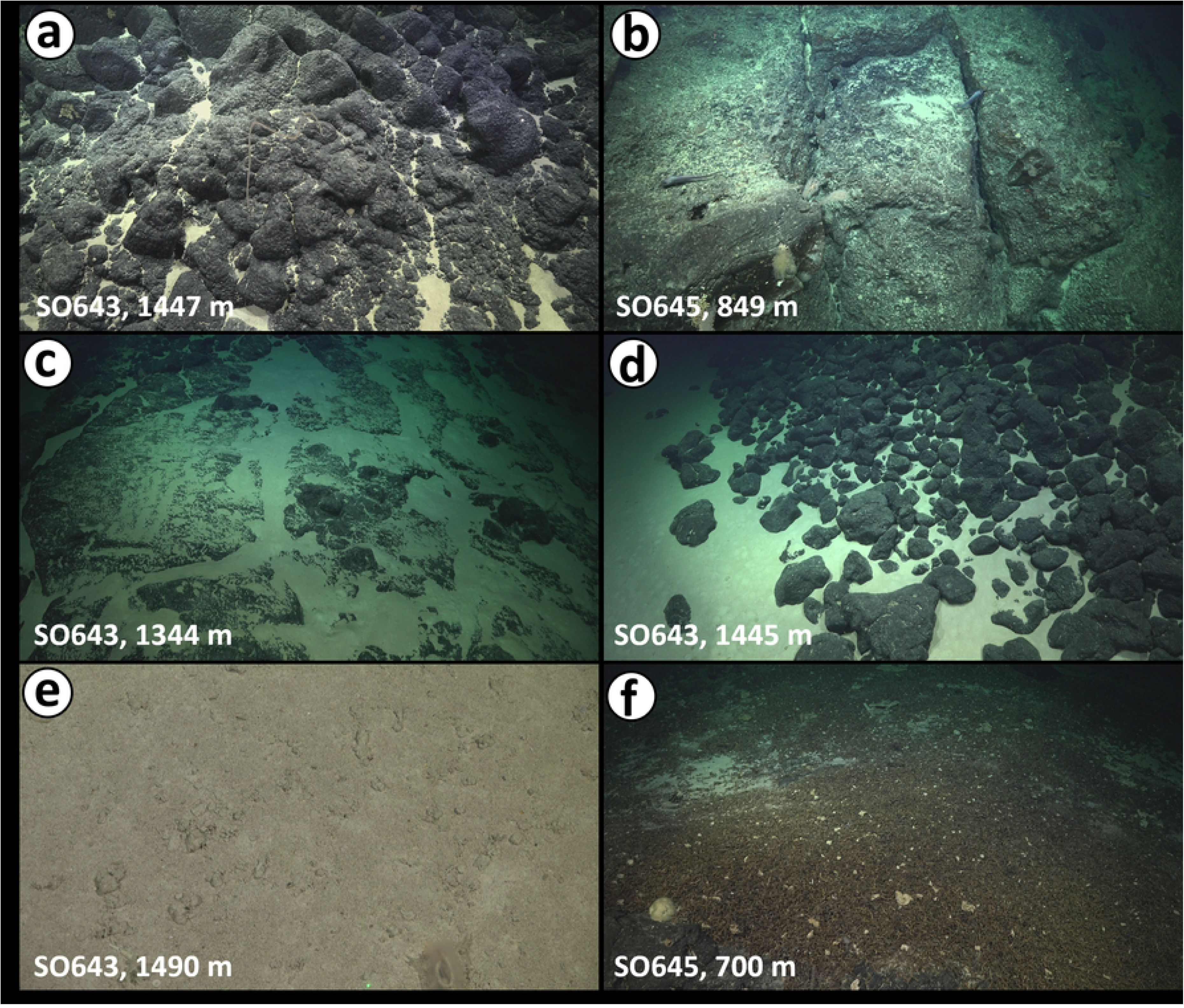
Examples of classified substrate types along the remotely operated vehicle (ROV) dive tracks. (a) and (b) are bedrock, (c) bedrock with sediment veneer, (d) sediment–rock transition, (e) sediment, and (f) coral rubble.

**Figure 6.**
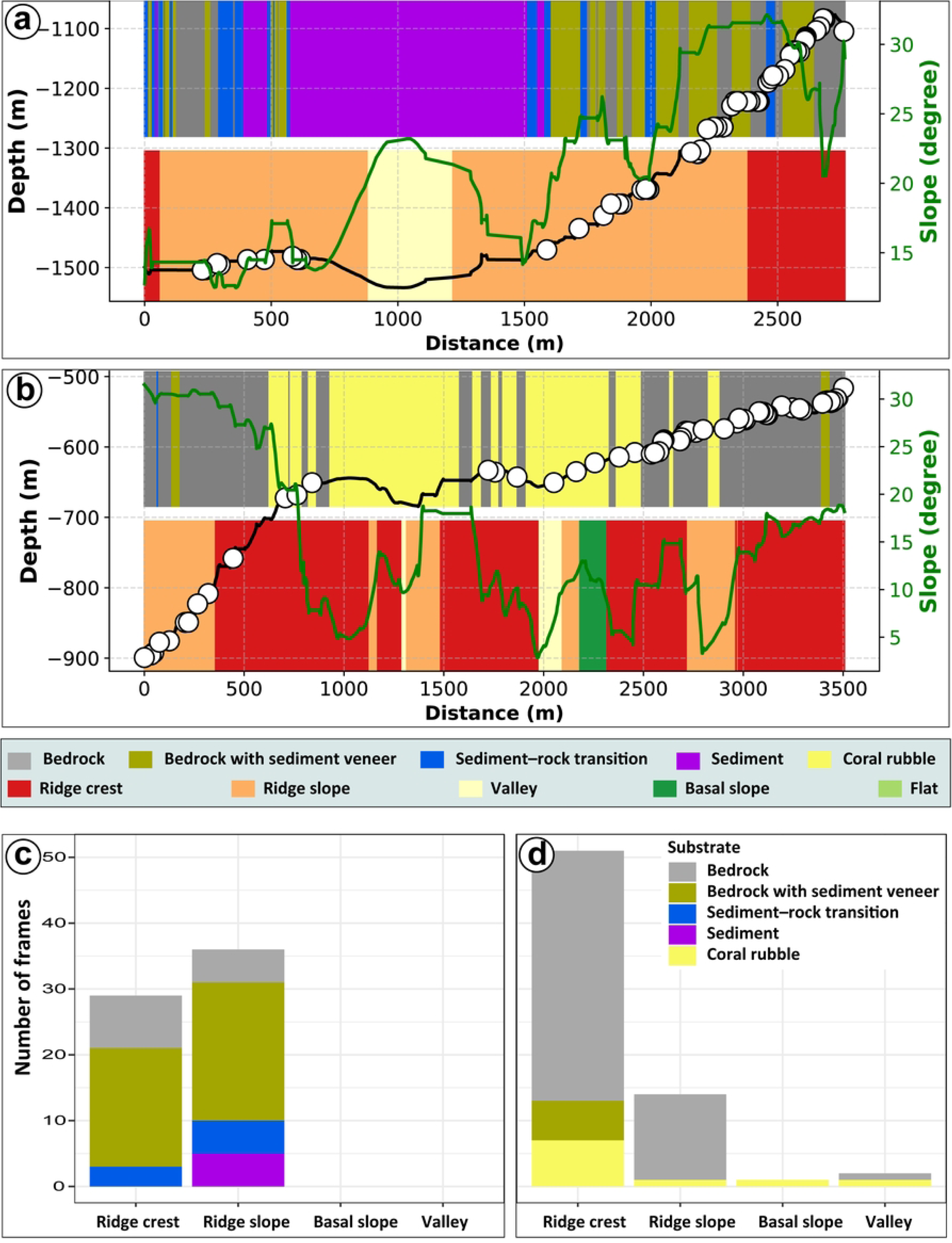
Spatial visualisation of quantitative megahabitat and substrate types identified on Solito Seamount. Depth (black line) and slope (green line) profiles along the two remotely operated vehicle (ROV) dive tracks with megahabitats (lower half) and substrate types (upper half) for (a) dive SO643 and (b) dive SO645. Depth profile showing only locations where the seafloor was visible and clearly resolved in the ROV video footage. White circles represent biological observations. Number of frames of substrate types observed in different habitats along ROV dive SO643 (c) and dive SO645 (d).

**Table 2.** Substrate types, their dominant substrate composition, structural complexity and sediment coverage observed along the ROV dive tracks of the Solito Seamount.

| Substrate type | Description | Dominant substrate composition | Structural complexity | Sediment coverage |
| --- | --- | --- | --- | --- |
| Bedrock | Massive volcanic outcrops or crusts forming hard, rugged surfaces with little or no sediment accumulation. | Lava flows, pillow lavas, boulders or cobbles | High | Absent (<5%) |
| Bedrock with sediment veneer | Volcanic bedrock outcrops partially draped by a thin veneer of sand or silt; sediment accumulates in cracks or depressions. | Lava flows, boulders or cobbles | High | Thin (5–20%) |
| Sediment–rock transition | Mixed mosaic of exposed volcanic rock, cobbles, and interstitial sediments; exhibit high occurrence of small-scale heterogeneity. | Sediments and bedrocks or cobbles | Moderate to high | Variable (10–50%) |
| Sediment | Uniformly distributed unconsolidated soft sediment forming a smooth or rippled seabed with low relief; no visible bedrock. | Unconsolidated sediments, mostly coarse grained | Low | High (>80%) |
| Coral rubble | Accumulations of dead coral fragments, skeletal debris, and shells, often mixed with sediment and occasionally overlying hard substrate. | Biogenic sediments | Low to moderate | Variable (20–50%) |

Across both dives, substrate distribution closely corresponded with seafloor habitats (Figs. 4 and 6). Ridge crests and ridge slopes were consistently dominated by hard substrates (bedrock and bedrock with sediment veneer), reflecting minimal sediment accumulation, whereas valleys and basal slopes favoured softer sedimentary substrates with lower structural complexity (Fig. 6). At SO643, ridge crest habitats were dominated by bedrock with sediment veneer (∼60–65%), with exposed bedrock (∼25–30%) and minor sediment–rock transition (∼10%). Ridge slopes were more heterogeneous, with bedrock with sediment veneer remaining dominant (∼55–60%) and exposed bedrock, sediment–rock transition, and sediment each contributing ∼10–15%. The valley megahabitat was characterized by smooth, low-relief seafloor composed mainly of unconsolidated sediments (Fig. 6a). At SO645, ridge crest and ridge slope were overwhelmingly composed of exposed bedrock (∼70–90%), while the coral rubble occurred locally (∼10–15%). The basal slope megahabitat was also sparsely represented and consisted mainly of coral rubble.

### 3.3 Benthic taxa and multi-level habitat associations

#### 3.2.1 PERMANOVA

Predominant benthic taxa observed along the dive transects are listed in Table 3. Results of the PERMANOVA demonstrated that dive, megahabitat (nested within dive), and substrate (nested within megahabitat and dive) all contribute significantly to variation in benthic community structure. A significant effect of dive was detected (Pseudo-F = 2.02, p = 0.008; S3 Table), likely reflecting unmeasured variation between dives, such as depth or flow differences. Megahabitat nested within dive explained additional variation (Pseudo-F = 1.64, p = 0.001; S3 Table), indicating that different geomorphological megahabitat classes within a dive contain different assemblages. The effect of substrate nested within megahabitat was strongest (Pseudo-F = 2.19, p = 0.001; Fig. 8, S3 Table) and contributed the most to the variance explained, indicating the largest structuring influence on benthic assemblages. Pairwise comparisons of megahabitat within each dive show that megahabitat effects were not consistent across dives. In SO643, benthic assemblages differed significantly between ridge crest and ridge slope habitats (t = 1.40, p = 0.005; Fig. 8, S4 Table). In contrast, no megahabitat pairs were significantly different within SO645, although several comparisons could not be tested due to insufficient replication (S4 Table). Pairwise tests of substrate nested within megahabitat and dive confirmed strong but variable substrate effects. In SO643, significant differences were detected among several substrate types within the ridge slope, with sediment-associated communities being consistently different from those on other substrate types (p < 0.01) (S5 Table). Weaker differences between substrate types were found in the ridge crest in the same dive. In SO465, conversely, substrate differences were most pronounced within the ridge crest, where coral rubble differed significantly from both bedrock and sediment veneer. Limited replication meant not all combinations could be tested.

**Table 3.** Dominant benthic organisms observed along the ROV dive transects of the Solito Seamount (for full descriptions of the benthic community see [97 in review]). Numeric label refers to their representation of SIMPER contributions in Fig 7.

| Numeric Label | Phylum | Class | Order | Family | ID |
| --- | --- | --- | --- | --- | --- |
| 1 | Porifera | Demospongiae | Tetractinellida | Ancorinidae | *Stelletta* sp. |
| 2 | Echinodermata | Echinoidea | Pedinoida | Pedinidae | *Caenopedina* sp. |
| 3 | Echinodermata | Echinoidea | Aspidodiadematoidea | Aspidodiademataidae | *Aspidodiadema* sp. |
| 4 | Echinodermata | Crinoidea | Comatulida | Antedonidae | Antedonidae sp. |
| 5 | Cnidaria | Octocorallia | Stolonifera | Alcyoniidae | *Anothothela* sp. |
| 6 | Cnidaria | Hexacorallia | Scleractinia | Dendrophylliidae | *Enallopsammia rostrata* |
| 7 | Cnidaria | Hexacorallia | Scleractinia | Caryophylliidae | *Madrepora oculata* |
| 8 | Cnidaria | Hexacorallia | Scleractinia | Caryophylliidae | *Caryophyllia* sp. |
| 9 | Cnidaria | Hexacorallia | Antipatharia | Schizopathidae | *Telepathes* sp. |
| 10 | Cnidaria | Hexacorallia | Antipatharia | Schizopathidae | *Bathypathes* sp. |
| 11 | Cnidaria | Octocorallia | Scleralcyonacea | Primnoidae | *Calyptraphora* sp. |
| 12 | Cnidaria | Octocorallia | Scleralcyonacea | Plexauridae | Plexauridae spp. |
| 13 | Cnidaria | Octocorallia | Scleralcyonacea | Keratoisididae | *Acanella* sp. |
| 14 | Cnidaria | Octocorallia | Scleralcyonacea | Coralliidae | *Hemicorallium* sp. |
| 15 | Cnidaria | Octocorallia | Scleralcyonacea | Chrysogorgiidae | *Chrysogorgia* sp. |
| 16 | Cnidaria | Hexacorallia | Actiniaria | Actinoscyphiidae | *Actinoscyphia* sp. |
| 17 | Arthropoda | Malacostraca | Decapoda | Nematocarcinidae | *Nematocarcinus* sp. |

The number of taxa contributing significantly to compositional differences differed between dives. For dive SO643 SIMPER identified a broad suite of taxa contributing to habitat, substrate and combined habitat-substrate contrasts, including reef-building scleractinians, octocorals, antipatharians, actiniarians, echinoderms and sponges (Fig. 7). In contrast, dive SO645 exhibited fewer taxa that contributed significantly to assemblage dissimilarities. The main contributing taxa were the sponge *Stelletta* sp, the gorgonians *Chrysogorgia* sp. and Plexauridae spp., and the shrimp *Nematocarcinus* sp., often with one or two taxa accounting for the majority of the dissimilarity.

**Figure 7.**
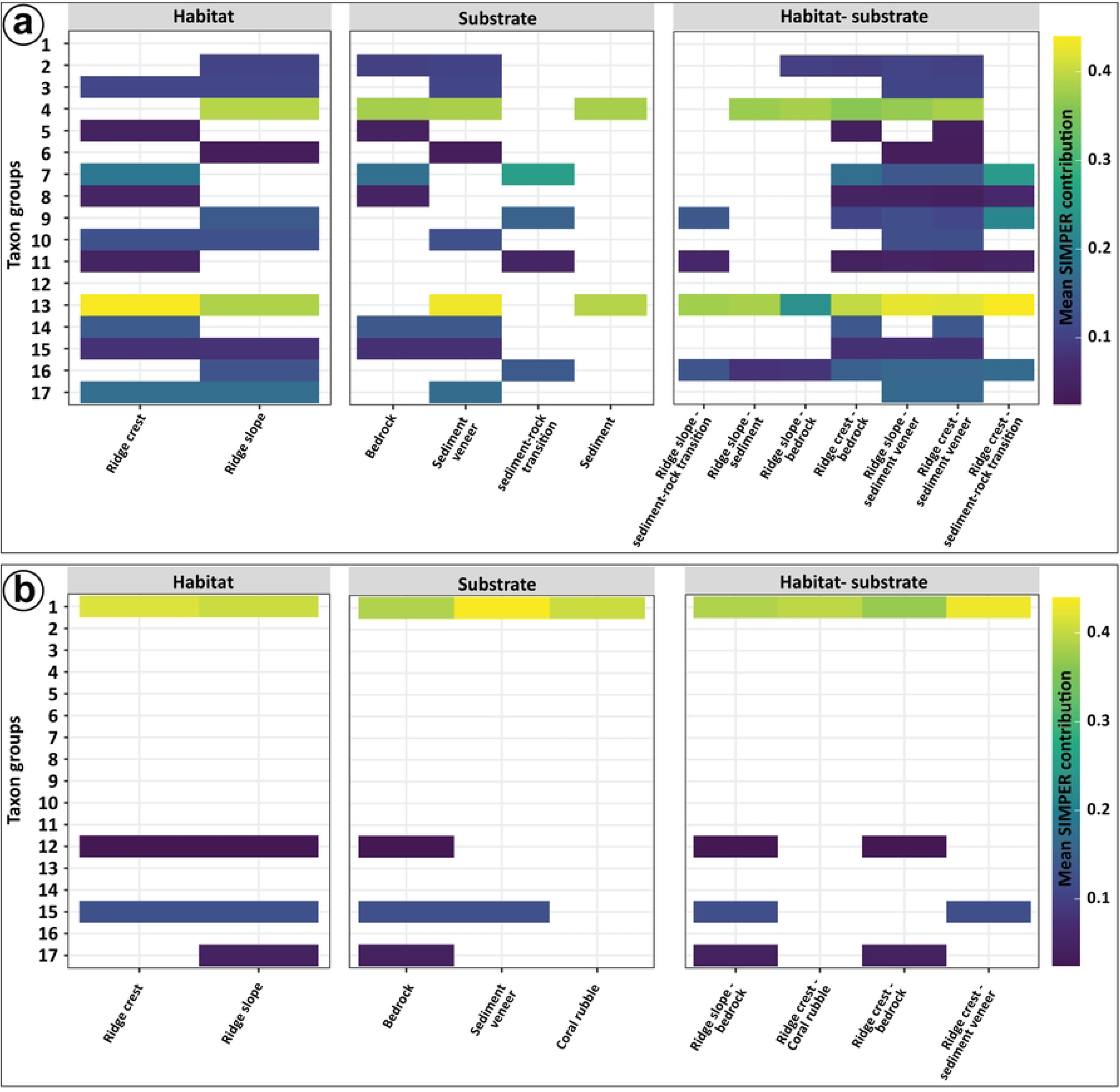
Distribution of faunal groups within the different habitats (left), substrates (middle) and combination of habitats and substrates (right), along remotely operated vehicle dive (a) SO643 and (b) SO645. Taxa groups are displayed in Table 3.

#### 3.3.2 Megahabitat associations (megahabitat-only SIMPER)

When taxa associations were assessed per megahabitat (Fig. 7), several topography-linked patterns were evident, being the most pronounced in dive SO643. Dissimilarities among S0643 ridge crest and all other megahabitats were predominated by reef-building and erect suspension feeders, including *Solenosmilia* sp.*, Caryophyllia* sp.*, Enallopsammia* sp.*, Calyptrophora* sp.*, Chrysogorgia* sp. and *Hemicorallium* sp. This predominance reflects a distinct summit assemblage characterized by hard-bottom reef-forming and branching taxa. The S0643 ridge slope megahabitat supported a mixed assemblage of antipatharians (*Telopathes* sp.), gorgonians (*Acanella* sp.), actiniaria (*Actinoscyphia* sp.), and crinoids (Antedonidae sp.). Associations for S0643 basal slope and valley megahabitats could not be reliably resolved due to low replication. In dive SO645, megahabitat similarity is higher than in SO643 with a more restricted set of taxa showed meaningful megahabitat associations. Octocorals Plexauridae sp., *Chrysogorgia* sp., and the sponge *Stelletta* sp. were representative of S0645 ridge crest and ridge slope habitats. Nematocarcinid shrimp (*Nematocarcinus* sp.) differentiated S0645 ridge crest and ridge slope habitats, even where substrate types were identical, suggesting that the geomorphic setting can influence faunal structure independently of substrate in SO645.

#### 3.3.3 Macro-scale substrate associations (substrate-only SIMPER)

When substrate type was examined independently within each dive, substrate effects were strong in both dives but most clearly resolved in dive SO643 (Figs. 7 and 8). Hard substrates hosted typical hard-bottom suspension feeding taxa that depend on hard surfaces for attachment. Bedrock and bedrock with sediment veneer substrates both had high contributions from crinoids (Antedonidae), the corals *Hemicorallium* sp., and *Chrysogorgia* sp. and urchins *Caenopedina* sp. (Fig. 8). Some of these taxa were only associated with outcropping bedrock (*Solenosmilia* sp.*, Caryophyllia* sp., Stolonifera sp.) or areas of bedrock with sediment veneers (*Acanella* sp.*, Bathypathes* sp.*, Nematocarcinus* sp.*, Aspidodiadema* sp.), showing some differentiation in affinities for hard or semi-hard substrate. Areas of mixed (sediment–rock transition) substrate hosted primnoid corals *Calyptrophora* sp., actiniarians (*Actinioscyphia* sp.), and black corals (*Telopathes* sp.) but also some reef-building corals (*Solenosmilia* sp.). Sedimented areas did not support a distinct fauna, although crinoids (Antedonidae) and bamboo-corals *Acanella* sp. were also found associated with these soft substrates (Fig. 8). In dive SO645, fewer taxa were resolved with distinct substrate signals. Areas of bedrock hosted gorgonians (Plexauridae spp.) and nematocarcinid shrimp, which contributed to contrasts across megahabitat types. Bedrock with sediment veneer was distinguished mainly by *Chrysogorgia* sp. reflecting a shift in community composition with an increase observed coincident with sediment-veneered surfaces. The clearest substrate signal was associated with coral rubble. Across all substrate pairs involving coral rubble, the sponge *Stelletta* sp. showed the highest contributions, indicating a tight association with coral rubble. Whilst *Stelletta* sp. occurred on other substrates, its predominance on areas of coral rubble substrates shows a distinct sponge-assemblage association.

**Figure 8.**
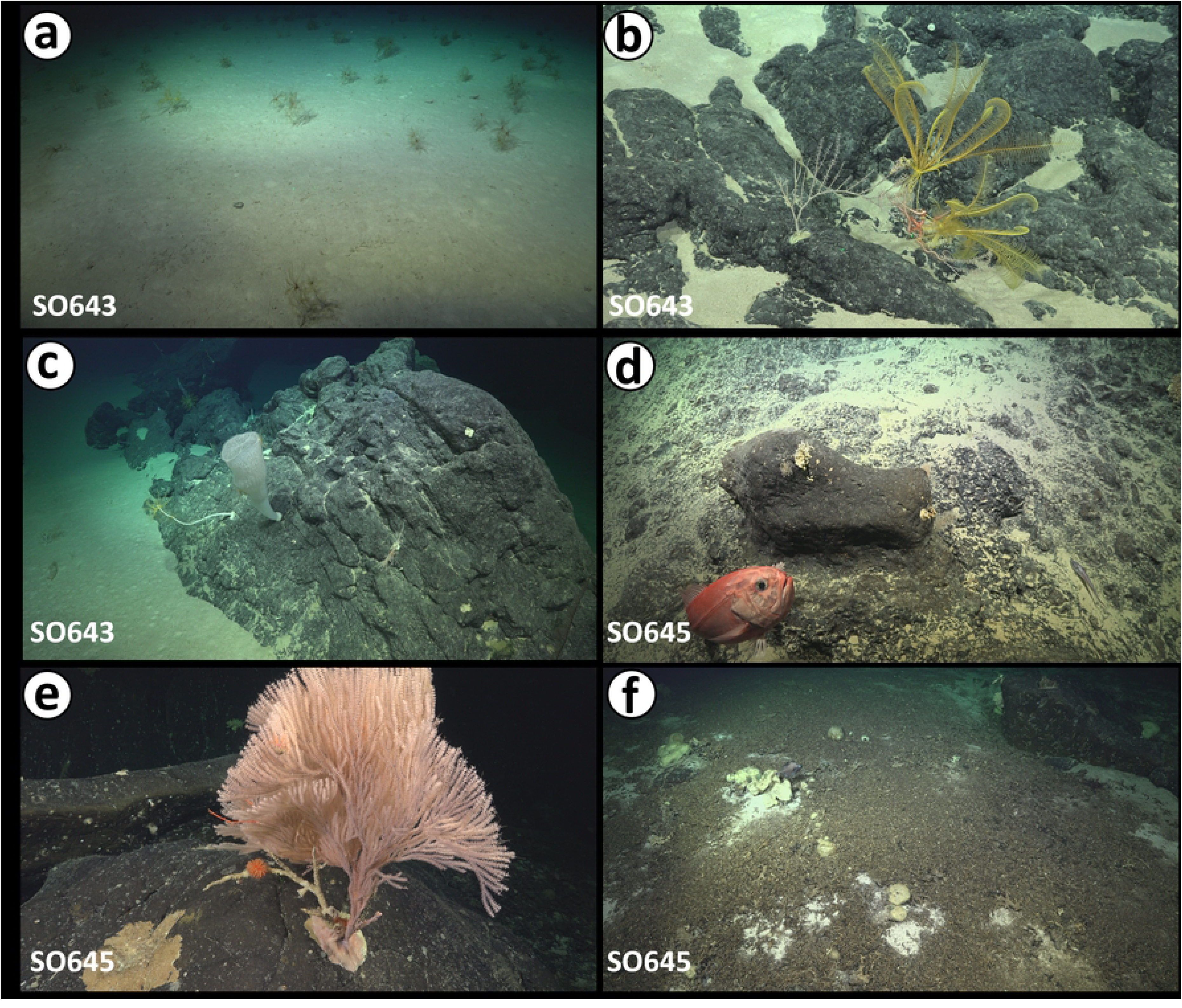
Representative examples of megafauna observed on different substrate types. (a) Fields of small bamboo corals (*Acanella* sp.) and an antedonid crinoid on soft sediment. (b) Antedonid crinoid on a sediment–rock transition zone. (c) Stony coral *Ellanopsammia rostrata* on bedrock within a sediment–rock transition zone, occasionally used as habitat by the orange roughy (*Hoplostethus atlanticus*). (d) Euplectellid glass sponge (*Regadrella* sp.) attached to bedrock partially covered by a thin sediment veneer. (e) Large branched primnoid corals (*Calyptrophora* sp.) on exposed bedrock. (f) Sponge field (Stelletta sp.) colonizing coral rubble derived from *Madrepora oculata*.

#### 3.3.4 Combined megahabitat-substrate associations

The combined megahabitat-substrate SIMPER (Figs. 7 and 8) clarified how megahabitat and substrate type interact to shape benthic assemblages. Overall, the combined analyses show that substrate modifies habitat level patterns, but the relative importance of these controls differed with depth and setting. In the deeper dive SO643, community structure is primarily organised by megahabitat type, with substrate-driven differences within habitat. In the shallower dive SO645, substrate type exerts a strong influence on assemblage composition across megahabitats, with habitat-level effects. The number of taxa contributing significantly to the SIMPER also contrasted between dives, with SO643 consistently exhibiting a broader suite of contributing taxa than SO645.

In SO643, the most distinct assemblages occurred in the ridge crest (bedrock with sediment veneer, followed by bedrock and sediment–rock transition). These ridge crest assemblages were consistently characterised by high contributions from reef-building corals (*Solenosmilia* sp.*, Enallopsammia* sp.), solitary corals (*Caryophyllia* sp.), octocorals (*Calyptrophora* sp.*, Chrysogorgia* sp.), red coral (*Hemicorallium* sp.), and malacalcyonacea (Malacalcyonacea sp.) (Fig. 8). Within the ridge crest megahabitat changes in substrate type correspond to changes in the relative contributions of framework-forming and branching suspension feeders. *Solenosmilia* sp. is a key discriminator between bedrock and sediment–rock transition or bedrock with sediment veneer. Solitary corals, primoids *Calyptrophora* sp. and *Chrysogorgia* sp. also contribute to the bedrock contrast, as do *Hemicorallium* sp. and Malacalcyonacea sp. On the S0643 ridge slope megahabitat, contrasts among substrates were driven primarily by large suspension feeders and echinoderms. Substrates comprising bedrock with sediment veneer hosted the most diverse assemblage, followed by sediment–rock transition and finally areas of sediment. Whilst the majority of taxa observed on the ridge slope were linked to bedrock with sediment veneer, black coral *Telopathes* sp., primnoid *Calypthrophora* sp., octocoral *Acanella* sp. and actiniarian *Actinoscyphia* sp. were found across multiple substrate types (Fig. 8). In SO645, the combined habitat-substrate patterns were simpler but still revealed a few strong signals. In the S0645 ridge crest megahabitat, coral rubble was a uniquely structured habitat, reinforcing that coral rubble represents a distinct macro-habitat supporting a distinct assemblage dominated by *Stelletta* sp. Differences between hard substrate macrohabitats (bedrock vs bedrock with sediment veneer) in the ridge crest areas were mainly driven by *Chrysogorgia* sp. which consistently had higher contributions in sediment veneer habitats. In the S0645 ridge slope megahabitat however, bedrock was characterised by Plexauridae and *Nematocarcinus* sp., differentiating slope from crest.

## 4. Discussion

### 4.1 Automated quantitative and multi-scale megahabitat mapping

Deep-sea habitats are inherently hierarchical, with physical and biological processes operating across mega-, macro- and micro-scales that shape species distributions, community composition, and ecosystem functioning [51,52,55,95,107–109]. Seamounts in particular exhibit environmental heterogeneity due to variations in topography, geomorphology, substrate, and hydrodynamics, creating distinct habitats that influence benthic assemblages [20,26,33,48,66]. Despite the ecological importance of these multiscale patterns, application of quantitative and automated deep-sea benthic habitat mapping remains uncommon, largely constrained by limitations in data resolution, spatial coverage, and methodological standardization. Previous unsupervised seamount benthic habitat mapping studies have often relied on bathymetric derivatives interpreted using subjectively defined thresholds or expert-driven classifications, limiting reproducibility and cross-site comparability [62,110]. In contrast, objective, data-driven approaches, which have been widely used in marine habitat mapping, offer the potential to objectively delineate habitat units across multiple spatial scales while explicitly capturing geomorphological complexity and environmental gradients [50,79,111–113].

In this study, the integration of multibeam bathymetry, seafloor terrain derivatives and backscatter intensity data enabled unsupervised characterization of the benthic habitats of the remote Solito Seamount across multiple scales. Metrics such as slope, curvature, BPI, and ruggedness captured both fine- and broad-scale geomorphological complexity, serving as proxies for habitat structure [50,56,79]. PCA combined with unsupervised k-means clustering enabled the segmentation of the seafloor into megahabitat classes defined by distinct combinations of depth, slope, rugosity, and substrate reflectance (backscatter intensity). Similar multivariate approaches have been applied in several deep-sea habitat mapping projects, particularly in studies of submarine canyons and seamount environments [63,65,66]. These classes delineate spatially contiguous geomorphological zones that likely represent ecologically meaningful benthic habitats (Fig. 4). Once integrated with visually validated ROV observations, these automated classifications enabled explicit links between geomorphology, substrate type, and benthic assemblages, reinforcing the ecological relevance of the mapped habitat units (Figs. 6 and 7).

While the K-means clustering provides a simple, reproducible framework for identifying seafloor classes, it is constrained by inherent methodological assumptions, often resulting in fragmented or spatially discontinuous habitat units [50,114]. In contrast, supervised or semi-supervised learning approaches such as Random Forest [115,116], Support Vector Machines [117,118], or more advanced convolutional neural networks [119–121] could generate more robust quantitative habitat maps. These methods incorporate ground-truth information providing stronger and more consistent integration of acoustic data with the substrate types and benthic assemblages. However, the two ROV transects in this study did not cover the full bathymetric range of Solito Seamount, limiting the availability of representative training data required for supervised classification. Such constraints are common in deep-sea studies, where the spatial coverage of ground-truth observations is often restricted. Consequently, k-means clustering was adopted as an objective, data-driven approach suitable for preliminary habitat characterization until more extensive surveys enable the development of supervised classification models. Future research should therefore prioritize the systematic collection of spatially representative biological and substrate observations to support supervised classification frameworks. This advancement would provide a stronger foundation for accurately representing seafloor heterogeneity, quantifying biodiversity patterns, and informing evidence-based marine spatial planning and conservation strategies.

### 4.2 Megahabitat and substrate on the Solito Seamount

The megahabitat structure identified via the K-means clustering approach for Solito is primarily controlled by depth, slope gradient, and surface roughness and hardness (backscatter intensity). The organisation of the five megahabitats aligns closely with the morphology of a cone-shaped seamount, where ridge and valley systems radiate downslope from a relatively narrow summit, and steeper volcanic slope, transitioning progressively into the surrounding abyssal plain (Fig. 4a) [10,122]. This configuration strongly influences substrate distribution. The limited summit area and higher relief restrict the accumulation of unconsolidated particles, resulting in widespread exposure of volcanic hard substrate around the summit area. Accordingly, ridge crests and ridge slopes are predominated by exposed bedrock and sediment veneer, with only localized sediment accumulation in small topographic depressions and fracture-controlled pockets (Figs. 5c and 5d). In contrast, sediment cover increases downslope as gradients decrease, with valleys acting as sediment accumulation zones where reduced slopes and topographic confinement favour sediment retention. This megahabitat and substrate organisation differs from the flat-topped, guyot-dominated seamounts of the NZ and S&G ridges, which are characterised by broad, flat summits, comparatively subdued relief, and laterally extensive sediment-covered platforms, resulting in proportionally less-developed slope environments [40,123].

Coral rubble represents a spatially limited but ecologically distinct substrate type on Solito, preferentially associated with shallower depths (dive SO645) and intermediate slopes (Figs 4 and 6). The coral rubble on Solito Seamount is derived primarily from framework-forming corals such as *Madrepora oculata* and bamboo corals, taxa widely recognized as ecosystem engineers that enhance seafloor heterogeneity and promote benthic diversity by creating a range of ecological niches [124,125]. Its occurrence across multiple megahabitats suggests that coral rubble distribution is not solely controlled by large-scale morphology, but rather reflects a combination of biological production, physical fragmentation of biogenic frameworks, and local hydrodynamic processes that govern downslope transport and retention [126–128]. As such, coral rubble may function as a transitional substrate between unconsolidated sediments and exposed bedrock [128–130], contributing additional structural complexity on Solito.

### 4.3 Benthic fauna on the Solito Seamount

A from both ROV dives elucidate that the habitat complexity and substrate composition are primary drivers shaping benthic megafaunal assemblages [131–133]. In dive SO643, distinct differences in assemblage composition between ridge crest and slope were apparent, likely driven by taxa such as bamboo coral *Acanella* sp., which showed contrasting abundances among habitats, with higher abundances in ridge slope (Fig. 7). Similarly, *Antedonidae* crinoids reached their highest abundances in ridge slope habitats. *Acanella* sp. was more abundant in sediment rich environments rather than bedrock with sediment veneer settings (Fig. 7). This pattern is consistent with observations for the congener *Acanella arbuscula*, which can occupy different substrate types [134], but is more commonly associated with soft-sediments [135]. In contrast, Antedonidae crinoids exhibited higher abundances on bedrock substrates, followed by sediments and bedrock with sediment veneer (Fig. 8). Bamboo corals and crinoids are suspension feeders that rely on suspended organic particles and pelagic prey for their nutrition [e.g., 136,137]. The higher abundance of these taxa on ridge slopes may be linked with locally accelerated currents, which facilitate the transport and/or resuspension of organic particles and zooplankton, contributing to increase the abundance of certain taxa [138,139].

Across the shallower dive SO645, benthic assemblage structure did not show marked differences among habitats (Fig. 7). However, the ridge crest showed substantial variation of taxa among coral rubble, bedrocks, and bedrock with a sediment veneer. Changes in the abundance of the sponge *Stelletta* sp. were particularly important explaining the dissimilarity of benthic assemblages among habitats and substrate types, showing higher abundances on coral rubble, followed by areas of outcropping bedrock, with the lowest abundances associated with bedrock with sediment veneer (Fig. 8f). Sponges are predominant components of shallow coral rubble macrohabitats and are considered key organisms due to their role in rubble binding and stabilization processes, as well as their role in the energy transfer through food webs [128]. Moreover, several sponge species exhibit significant habitat preference for non-living substrates, preferring dead corals, unconsolidated coral rubble and calcium carbonate rocks [140]. In deep-sea environments, coral rubbles are known to enhance the metabolic activity of cold-water coral reefs, where diverse faunal (including sponges) and microbial communities recycle considerable amounts of particulate (phytodetritus) and dissolved organic matter [141]. Therefore, we expected that coral rubble located on the ridge crest megahabitat constitutes a preferred habitat for *Stelletta* sp., associated with the broad availability of suitable microhabitat for settlement and the increased availability of organic matter sources. Comparable nursery functions associated with coral frameworks have been reported from seamount JF6 and other seamounts of the Nazca–Desventuradas region, suggesting recurrent functionality roles across biogeographically connected systems [43,44].

The structurally complex megafaunal benthic communities observed on Solito differed between dives in terms of composition and the number of observed taxa contributing to community differentiation. The deeper assemblages (SO643) are characterised by a comparatively diverse suite of megafaunal taxa, including reef-building scleractinians (*Solenosmilia* sp.*, Caryophyllia* sp.*, Enallopsammia* sp.), octocorals (*Acanella* sp.*, Calyptrophora* sp.*, Chrysogorgia* sp.), antipatharians, actiniarians (*Actinoscyphia* sp.), and echinoderms (Fig. 7). The shallower assemblage (SO645) was dominated by a smaller number of taxa with community differences primarily driven by a few abundant or structurally dominant species (*Stelletta* sp.). As the terrain metrics used in this study captured substantial habitat variability of the seamount, with the first two principal components explaining a large fraction of variation and over 90% captured by the first five PCs (S2 Fig), additional variation in habitat structure could be explained by unquantified factors, such as oxygen concentration and current regimes. The shallower assemblages observed on dive SO645 coincide with the ESSW (∼50–600 m) above the lower boundary of the OMZ, whereas the deeper communities in dive SO643 occur within well-oxygenated AAIW (600–1,300 m) and PDW (∼1,500–3,000 m) [72]. OMZs strongly shape the characteristics of both pelagic and benthic populations [142], and enhanced richness is often observed below the OMZ boundary, where reduced oxygen limitation and increased organic matter availability support more diverse benthic assemblages[143,144]. Consistent with this pattern, complementary analyses showed variation in species composition on Solito seamount mainly driven by variations in salinity, depth, temperature, and dissolved oxygen (Tapia-Guerra et al. in review). Overall, the benthic megafaunal assemblages on Solito exhibit both depth-related environmental gradients and distribution of different substrate types. The presence of shared habitat-forming taxa and commercially important species with the Nazca-Desventuradas and Juan Fernández systems, suggests that Solito may contribute to biogeographic stepping stone within the southeastern Pacific seamount network, potentially facilitating faunal connectivity across adjacent area (Gorny et al., 2017; National Geographic et al., 2017; Tapia-Guerra et al., 2021b). The quantitative baseline provided here highlights Solito as a priority candidate for inclusion in regional marine spatial planning and conservation strategies, particularly given its habitat heterogeneity and likely role in sustaining deep-sea connectivity.

## 5. Conclusion

This study demonstrates the value of integrating automated objective multi-scale megahabitat mapping with *in situ* biological observations to resolve the physical and ecological organisation of a deep-sea seamount. By combining high-resolution multibeam bathymetry and backscatter intensity data with ROV-based observations of substrate and megafauna, we developed a quantitative and reproducible framework that links geomorphology, habitat structure, and benthic community patterns at Solito. The automated, multi-scale approach advances beyond traditional rule-based methods, provides the physical context necessary to interpret variation in megafaunal composition, abundance, and biodiversity. Solito supports diverse and structurally complex benthic communities dominated by cnidarians, echinoderms, sponges, and arthropods. The distribution and composition of benthic megafauna on Solito are strongly influenced by megahabitat type and substrate heterogeneity. Community differences between dives highlight the interaction of habitat complexity and broader-scale depth associated environmental gradients in shaping assemblages below the oxygen minimum zone. With spatial limitations of ROV surveys, this study outlines a pathway toward more robust, and transferable deep-sea habitat mapping approach. More broadly, integrative frameworks such as the one presented here provide essential baselines for biodiversity assessment, spatial planning, and the conservation of vulnerable seamount ecosystems, particularly in regions facing increasing anthropogenic pressure and limited biological knowledge.

## CRediT authorship contribution statement

Conceptualization: YN, DJBS.

Formal analysis: YN, DJBS, JMT.

Investigation: YN, DJBS, JMT.

Methodology: YN, DJBS.

Resources: JS, EEE, HAS, AJJ.

Supervision: JS, EEE, HAS, AJJ.

Writing – original draft: YN, DJBS, JMT, GZH.

Writing – review & editing: YN, DJBS, JMT, GZH, JS, EEE, HAS, AJJ.

## Acknowledgements

We would like to thank the captain and crew R/V *Falkor (too)* (Schmidt Ocean Institute cruise FKt240108), and the onboard scientific personnel participating in and leading the expedition. We are grateful to the Juan Fernández community, Community Functional Organization (OCF) Mar de Juan Fernández, for their collaboration and support during the expedition. This research was funded by ANID FONDECYT projects 1241386 and ATE22004 (BiodUCCT), with additional support from Conservation International on behalf of Blue Nature Alliance (CI-115211), and The Nippon Foundation-Nekton Ocean Census Programme. JMT acknowledges ANID Subdirección de Capital Humano Doctorado Nacional 2022-21220674. GZH was funded by BECAS CHILE-Postdoctoral abroad program (ANID) and additional support was provided by Marine Symbiomes Research Group, Stazione Zoologica Anton Dohrn (SZN). The analysis and writing of this study were supported by the Minderoo-UWA Deep-Sea Research Centre, funded by the Minderoo Foundation.

## Figure captions

S1 Figure. Bathymetry and terrain derivatives used to map the megahabitats of the Solito Seamount. (a) Depth, (b) Backscatter intensity, (c) Slope, (d) Broad-scale BPI (bathymetric position index), (e) Fine-scale BPI, (f) Planform curvature (g) Profile curvature, (h) TRI (terrain ruggedness index) (i) VRM (vector ruggedness measure).

S2 Figure. Scree plot of principal component analysis (PCA) showing the proportion of variance explained by individual components (green bars) and the cumulative variance explained (blue line), with a 90% cumulative variance threshold indicated

S3 Figure. Violin plots showing the distribution of standardized (Z-score) terrain variables across megahabitat classes. BPI: bathymetric position index; TRI: terrain ruggedness index; VRM: vector ruggedness measure. Each violin represents the density distribution of values within a megahabitat class, with an embedded boxplot. In the boxplots, the centre line indicates the median, the lower and upper box boundaries represent the first and third quartiles (interquartile range), and the whiskers extend to the most extreme values within 1.5 × IQR. Points beyond the whiskers denote outliers.

